# Reagentless Biomolecular Analysis Using a Nanoscale Molecular Pendulum

**DOI:** 10.1101/2020.04.02.020453

**Authors:** Jagotamoy Das, Surath Gomis, Jenise B. Chen, Hanie Yousefi, Sharif Ahmed, Alam Mahmud, Wendi Zhou, Edward H. Sargent, Shana O. Kelley

## Abstract

The ability to sense biological inputs using self-contained devices unreliant on external reagents or reporters would open countless opportunities to collect information about our health and environment. Currently, a very limited set of molecular inputs can be detected using this type of sensor format. The development of versatile reagentless sensors that could track molecular analytes in biological fluids remains an unmet need. Here, we describe a new universal sensing mechanism that is compatible with the analysis of proteins that are important physiological markers of stress, allergy, cardiovascular health, inflammation and cancer. The sensing mechanism we developed is based on the measurement of field-induced directional diffusion of a nanoscale molecular pendulum tethered to an electrode surface and the sensitivity of electron-transfer reaction kinetics to molecular size. Using time-resolved electrochemical measurements of diffusional motion, the presence of an analyte bound to a sensor complex can be continuously tracked in real time. We show that this sensing approach is compatible with making measurements in blood, saliva, urine, tears and sweat and that the sensors can collect data *in situ* in living animals. The sensor platform described enables a broad range of applications in personalized health monitoring.

## Introduction

Self-regenerating sensors that can dynamically and continuously detect biomolecular species in physiological systems remain an unmet need^1,2^. The ability to monitor protein biomarkers *in vivo* would provide a powerful tool for disease monitoring and treatment response^3–5^. A key requirement for this type of sensing application is a reagentless assay format where all required elements are incorporated into a self-contained sensor to allow for autonomous function.

The most robust sensors for reagentless, dynamic physiological monitoring at the molecular level rely on intrinsically redox-active chemistry or pathways that can be monitored at an electrode surface^5–12^. Indeed, electrochemical sensors for glucose and lactate are the dominant sensors used for the development of dynamic detection systems for monitoring in physiological systems because redox-active enzymes can be used to report on the presence of the target analytes in the absence of added reporters. Reagentless electrochemical sensors that are instead affinity-based and compatible with *in vivo* monitoring applications have been generated based on DNA aptamers that serve as recognition elements^13–18^. While powerful for the collection of pharmacokinetic data in living systems where analyte concentrations are high, aptamer-based sensors typically have low binding affinities that render them incompatible with many sensing applications.

We posited that by monitoring the kinetics of diffusion at the nanoscale it would be possible to monitor binding events using simple electrochemical measurements. With a tethered sensor displaying a target-specific antibody, we hypothesized that we could leverage dynamic phenomena to observe events that are affected by subtle changes in molecular composition, with the measurement of small changes in diffusional behavior used as a signature of specific biomolecular complexation events. To develop this capability we first investigated whether it would be possible to extract diffusional profiles of molecular complexes using a simple theoretical analysis exploring the motion of a tethered nanoscale sensor complex – modeled as a molecular pendulum - and then pursued experimental studies to identify specific binding interactions using current-based readout.

## Results and Discussion

To explore the diffusional behavior of biomolecular species tethered to an electrode surface, we first modeled the behavior of a nanoscale molecular pendulum (NMP) under the influence of an applied electric field to determine the time (τ) that would be required to bring a complex to the surface (Figure 1 and Supplementary Material, Figures S1-S6). The NMP is subject to a hydrodynamic drag force (F_d_) as well as to forces caused by Coulombic interactions (F_c_) with neighboring NMPs. The diffusion of large biomolecules is predicted to be slow (> 10^−6^ cm^2^/s) even in the presence of a tether that restricts the diffusional directionality^19^, thus we introduced a negatively-charged, rigid linker that would respond to a positive field generated with an applied potential. This electrostatic interaction introduces a force opposing the hydrodynamic drag that reflects the interaction with the electric field (F_e_).

**Figure 1.**
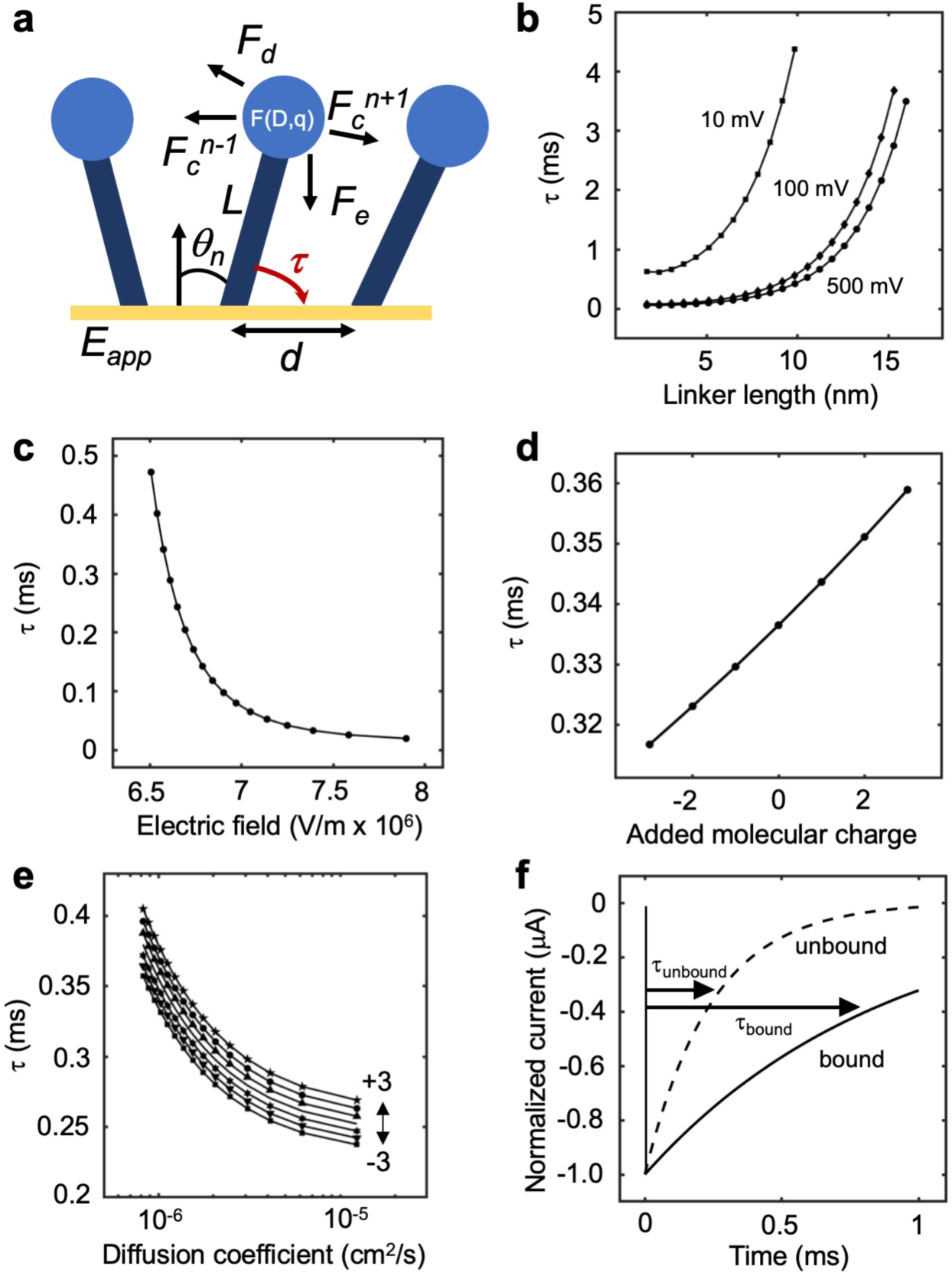
Modeling the dynamics of a nanoscale molecular pendulum (NMP) tethered to an electrode surface. **a) Model parameters.** The dynamics of the NMP were modelled by considering the drag force (F_d_), the force exerted by the applied field (F_e_), and electrostatic interactions between neighboring NMPs (F_c_). The length of the linker (L) separating the “bob” of the pendulum from the surface, the distance between neighboring NMPs (d), the average angle of the NMPs relative to the surface (θ), molecular charge (q), diffusion coefficient (D), and the applied electric field (E_app_) were all varied to explore NMP dynamics. Under an applied positive potential, a negatively charged NMP is attracted to the sensor surface. The transit time of the NMP is reflected in the modulation of τ. **b) Dependence of τ on the length of a negatively charged linker.** Linkers of at least 10 nm appear optimal for faster diffusional kinetics, with faster transit in higher fields. **c) Dependence of τ on the magnitude of the applied electric field.** As the average applied electric field increases, τ decreases as the NMP is attracted towards the electrode surface. This is expected for a negatively charged NMP accelerating through the field gradient of a positively applied potential. **d) Dependence of τ on added molecular charge.** Added molecular charge modulates the negative charge of the NMP and reduces or increases F_e_ to modulate τ. **e) Dependence of τ on diffusion coefficient**. As the diffusion coefficient of the analyte is increased with decreasing molecular weight and size, τ decreases as the effect of F_d_ is reduced. The addition of negative charges decreases τ further. **f) Modeled current decay** expected if the dynamics of the NMP could be monitored using an electrochemically-active label in the absence and presence of a molecular cargo 20 nm in size.

We explored (Figure 1b - f) the role of linker length, electric field strength, changes in molecular charge, as well as variations in cargo size and diffusion coefficient, and observed that all of these parameters modulated the τ values observed. With short linkers and weaker applied potentials, τ values approached 100 microseconds, whereas longer linkers that provide higher levels of charge in stronger fields exhibited smaller τ values approaching 20 microseconds. Small variations in molecular charge produced changes in τ, and these charge changes could be mapped on to variations in diffusional rates and coefficients to produce predictions for how τ would be affected by modification of the properties of the “bob” of the NMP. We then used the model to simulate electrochemical current transients as a function of a molecular size change of 20 nanometers. Significant changes to the current profiles were observed, indicating that changes in τ could be correlated to experimentally observable electrochemical currents with microampere resolution (Figure 1e).

We explored these phenomena experimentally by attaching a protein-binding antibody to a rigid DNA spacer (Figure 2, Figures S7-S10). Potential-dependent modulation of DNA orientation on electrode surfaces has been observed in previous studies^20–22^ and therefore this material appeared appropriate for the potential-responsive linker of the NMP. In order to obtain a redox reporter signal that could provide a means to monitor the dynamics of the NMP, ferrocene was attached to the DNA linker, which is oxidized at ∼ +400 mV versus Ag/AgCl. We therefore chose a potential of +500 mV to trigger the facilitated diffusion of the complex and the electrochemical reaction. As a modulator of the overall molecular weight and diffusional coefficient of the NMP, we selected the protein troponin I, which possesses a molecular weight of 24 kD. Attachment of a troponin-specific antibody to the DNA linker provided the needed specificity for the NMP.

**Figure 2.**
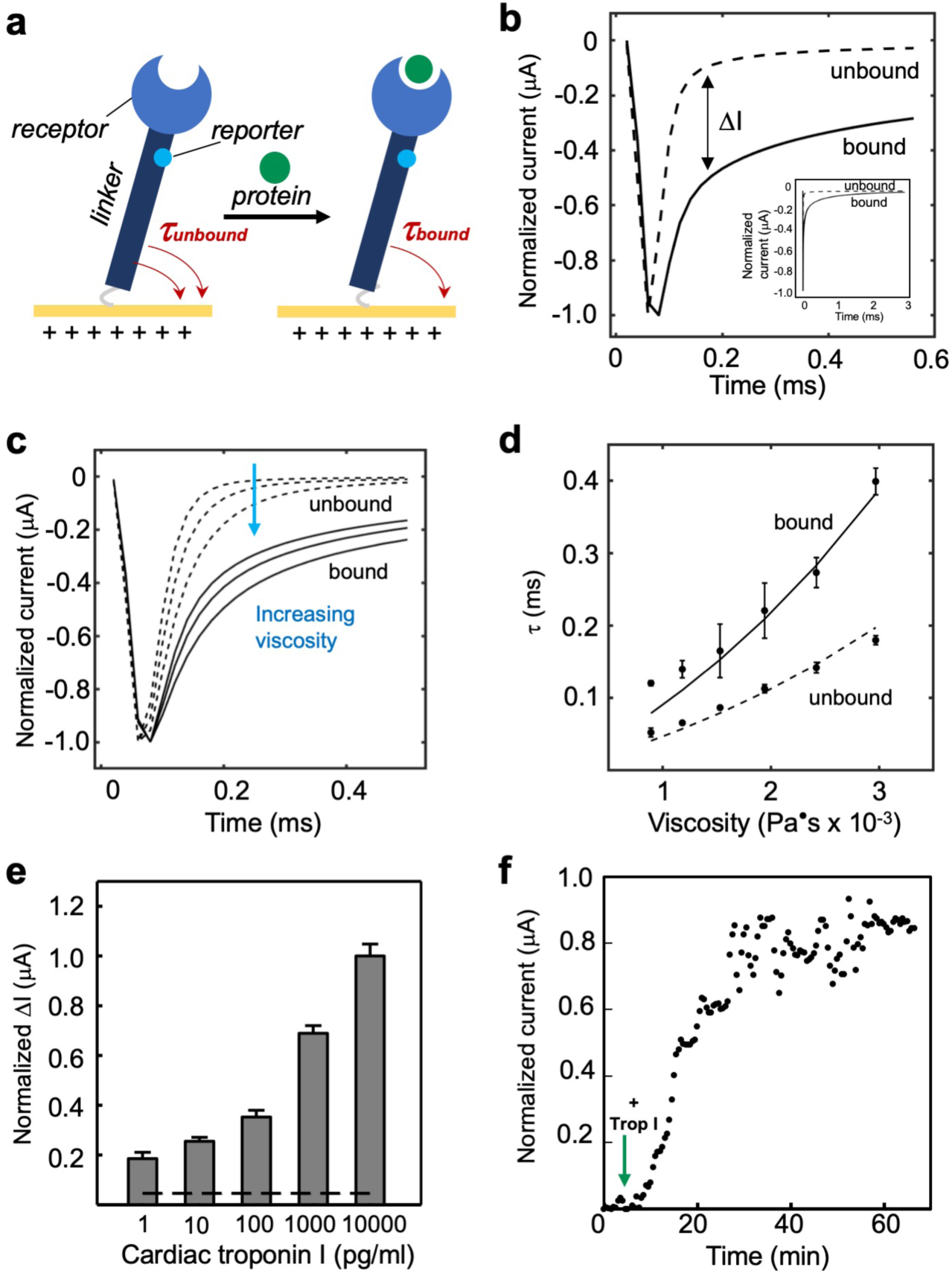
Modulation of NMP dynamics by protein binding. **a) A protein-binding NMP** was constructed using a DNA linker and an antibody specific for troponin I. A redox reporter, ferrocene, was incorporated into the DNA linker that would be oxidized at an electrode potential compatible with the electric field required to transport the NMP to the surface. **b) Observation of binding-induced modulation of NMP transport** using chronoamperometry in the presence (solid lines) and absence (dotted lines) of troponin I. The inset shows the full chronoamperometry traces collected with a potential step of +500 mV. Both τ and ΔI can be used to monitor the response of the system. **c) Dependence of NMP dynamics in bound and unbound states by modifying the fluid matrix.** By changing the viscosity of the medium to mimic different fluid conditions using increasingly higher glycerol concentrations, the NMP encounters higher drag force during transit and t increases. The bound sensor exhibits a larger τ value and thus slower current response. **d) Comparison of observed and calculated t values for NMPs in varied drag-modulated fluid matrices.** The theoretical approach (line plots) matches the decay characteristics of the experimental NMP time response (data points) for the bound and unbound states, allowing the characterization of NMP dynamics in varied fluid matrices. **e) Concentration-dependent signal change** for cardiac troponin I in buffered solution suggests that the bound state DI is distinguishable down to 1 pg/ml. Data from a control sensor has been subtracted. **f) Time-dependent current modulation for NMPs in the presence of troponin I** demonstrates the significant change in measurable current and equilibration within ∼ 20 minutes. The troponin I was injected into solution at t = 7 minutes.

Using chronoamperometry, we applied a potential and measured the faradaic current from the redox marker. For the NMP in its unbound state, a τ value of ∼100 microseconds was measured, whereas in the presence of added protein, the NMP in its bound state exhibited a τ value of ∼300 microseconds (Figure 2b). As a measured current difference at τ, the bound state exhibits upwards of 10 microamperes higher raw current compared to the unbound state. The experimental current transients also include contributions from capacitive charging, but the fast timescale of these events are not resolvable by our measurements which sampled current values every 20 microseconds.

To explore the variation in diffusional behavior of the tethered NMP, we varied the drag force on the tethered complex by altering the solution viscosity with added glycerol (Figure 2c). Calculation of the τ values as a function of the drag force investigated experimentally indicated good agreement with the model (Figure 2d), suggesting that we can calibrate our model across differing fluid matrices. We also observed increased τ values and resultant changes in current for the bound state of the NMP complex as a function of analyte (troponin) concentration (Figure 2e). At concentrations of troponin as low as 1 pg/ml (∼40 fM), statistically significant changes in current were observed. The increasing current differences as a function of protein concentration reflect the increasing levels of binding over the fM-pM concentration range. To explore the time dependence of the binding-modulated changes in diffusional behavior for the NMP, we collected current transients as a function of incubation time and determined that changes in the current could be observed within 10 minutes (Figure 2f). In the presence of non-target proteins or a sensor where the antibody was replaced by BSA, significant current modulations were not observed.

In order to challenge the specificity of the NMP for the target protein troponin I, we tested whether the concentration-dependent current changes would be retained in biological fluids including saliva, urine, tear fluid, blood and sweat (Figure 3a). Small fluctuations in the current transients, tabulated as ΔI values, were observed as a function of the type of fluid present; and statistically significant changes in current were observed with 1 pg/ml of troponin I for all sample types tested. Small variations in the magnitude of the effect were observed with the different sample types, likely due to changes in ionic strength or viscosity caused by the composition of the different fluids. The retention of performance in these different fluids is a key feature of this system that reflects the dynamic measurements that are the foundation of the approach.

**Figure 3.**
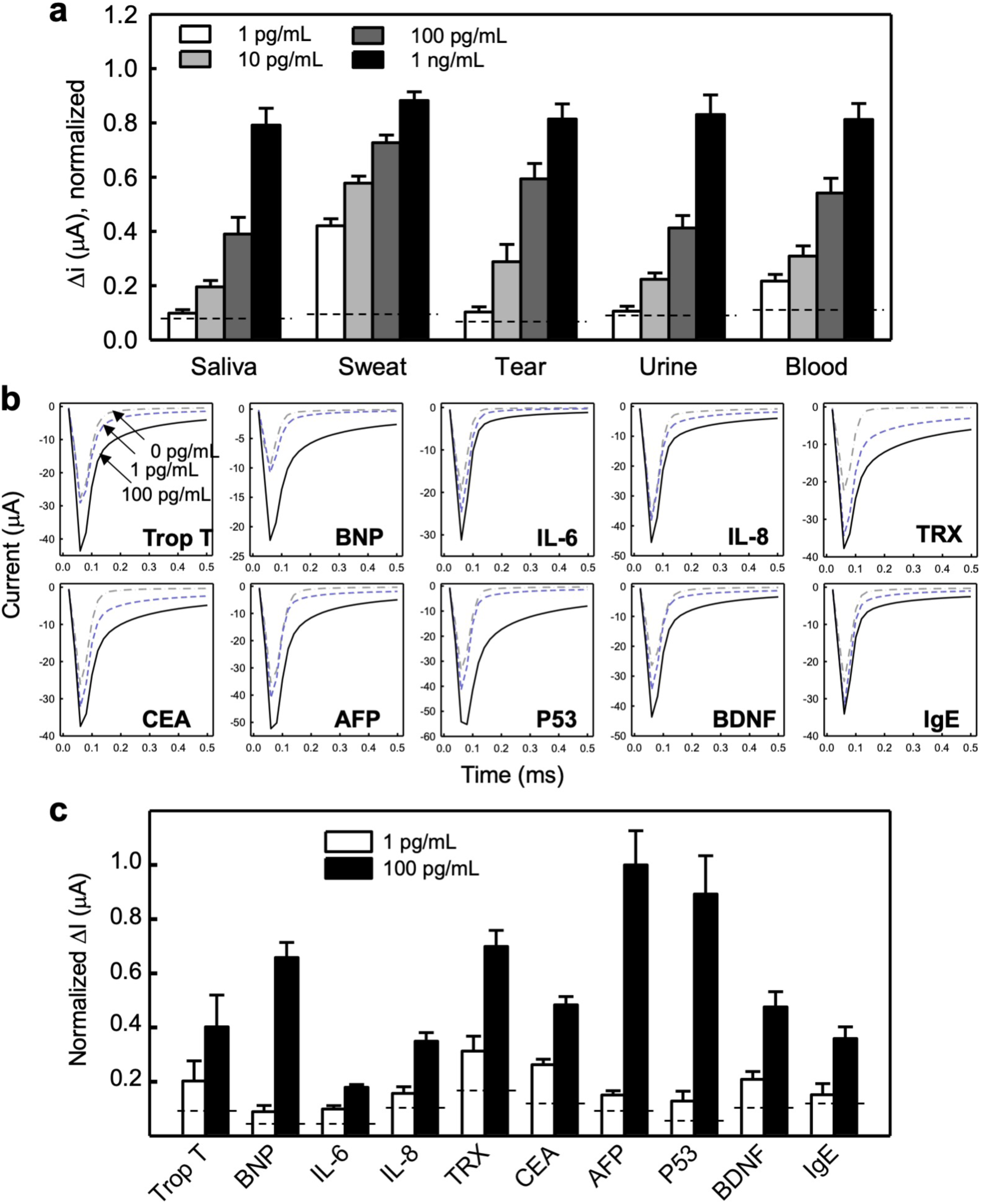
Panel of proteins and biofluids that can be monitored using NMPs. **a) Detection of cardiac troponin I in different biofluids** including saliva, sweat, tears, urine, and blood. Dotted lines represent cut-off values for limit-of-detection, including three standard deviations of values collected when NMPs are incubated with blank samples containing only the biofluid. **b) Testing of a panel of NMPs specific for cardiac, inflammation, stress, and cancer markers.** Changes in the current response curves are distinguishable between unbound and bound NMPs for each sensor tested. **c) Concentration-dependent responses for each protein tested**. Changes in the current response curves are distinguishable between unbound and bound NMPs.

A panel of proteins was tested to explore whether the binding-induced modulation of the NMP was a general effect (Figure 3c and S11). We tested a panel of 10 different proteins with varying charges, sizes, and molecular weights, using NMPs that featured antibodies specific for each protein. Chronoamperometry was used to monitor the change in τ values in the presence of two different concentrations of each protein, and each factor could be reproducibly monitored. Smaller proteins (e.g. IL-6) produced more subtle changes to the current transients for the corresponding NMPs, but still elicited statistically significant signal changes with τ variation down to ∼20 μs. These results demonstrate that diffusional behavior, when resolved using kinetic measurements, provides a universal reporter on biomolecular complexation status.

We then turned to exploring whether these constructs could be used for dynamic, continuous monitoring *in living systems* (Figure 4 and S12-14). We first tested the reversibility of the signals observed for troponin I-specific NMPs: here we observed that the current changes triggered by the presence of the protein disappeared within one hour when the protein was no longer present (Figure 4a). Multiple cycles of protein and blank incubation indicated that the NMPs were stable on the time scale of hours, and longer incubations between cycles showed that the system was stable over days (Figure 4a). NMPs could also be stored in biological fluids like saliva for multiple weeks and remained stable (Figure 4b).

**Figure 4.**
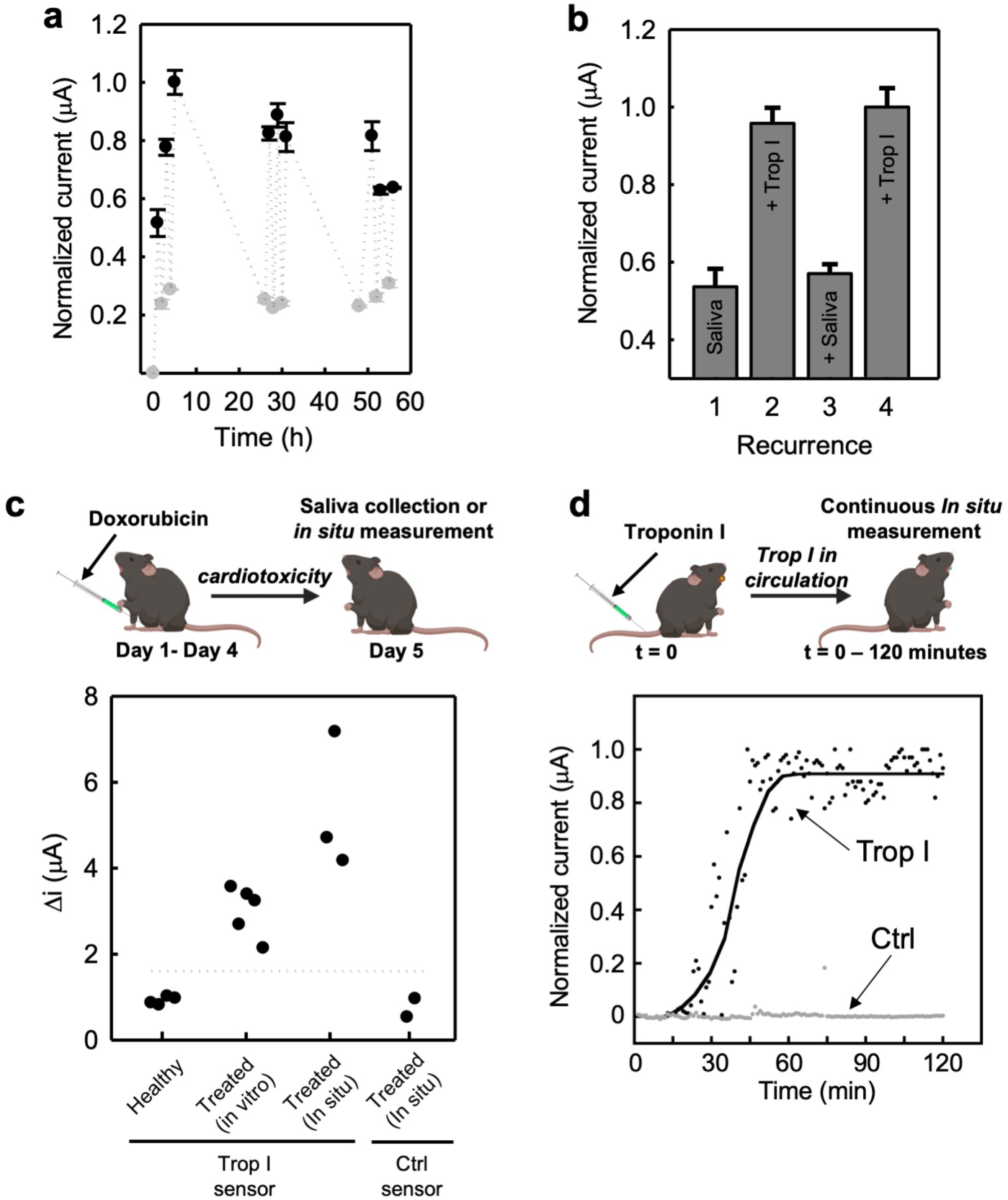
NMP-based monitoring of a cardiac marker in living animals. **a) Short-term cycling of NMP signals.** Black dots indicate signal change of the NMP when it is incubated with target protein (10 ng/mL) for 50 minutes and the gray dots represent signal change when the IMP is incubated with a buffered solution of a non-target protein (10 ng/mL) for at least 50 minutes. **b) Long-term incubation and response of NMP in saliva. NMP**s were stored in saliva for 3 weeks. After 3 weeks of incubation, NMP signals were obtained in saliva and saliva-spiked with troponin I (1 ng/mL) with 50 minute incubation cycles. **c) Measurement of troponin I in saliva both ex situ and in situ using the NMP.** To induce cardiac dysfunction, mice were treated with doxorubicin for four days and saliva samples were collected at day 5 for *ex situ/in vitro* measurements, or single point measurements were collected in situ within the mouth. **d) *In situ* continuous monitoring of cardiac troponin I in a murine model**. Mice were injected with cardiac troponin I 45 minutes before data acquisition and the NMP signal change was monitored continuously after 10 minutes of sensor equilibration with NMP featuring a troponin I antibody or BSA (ctrl).

We pursued the measurement of troponin I in living animals by testing two models. The first model was one that uses drug-induced cardiotoxicity to induce cardiac failure in mice^23^, which is known to induce the release of troponin into the blood stream and saliva (Figure 4c). Mice were treated for four days with doxorubicin, and on the fifth day the animals were tested in situ using NMP-functionalized electrodes (Figure S15 and S16). Elevated current levels were measured for a treated cohort of mice relative to untreated mice. Mice were also tested with control NMPs that did not possess a specific antibody and baseline levels of current were detected. The second model tested involved direct injection of troponin I into mice and continuous monitoring using a troponin I or control NMP implanted into the oral cavity of a mouse (Figure 4d). Within one hour after injection of the protein, significant increases in the differential currents were measured for the troponin I sensors, while the control sensor current levels remained at baseline.

These results indicate that sensors based on NMPs provide a general approach for the development of implantable sensors that can be used to monitor physiologically relevant proteins directly in biofluids. Given recent advances in the development of wearable monitoring devices^24–26^, this protein sensing concept may find broad applicability. Furthermore, the simple design of NMP sensors and compatibility with a wide range of targets including nucleic acids, antibodies and microorganisms the approach may also find utility in the development of clinical diagnostics. The development of versatile sensors that can track molecular analytes in biological fluids remains an unmet need now fulfilled by the performance exhibited by the NMP approach.

## Acknowledgements

### Funding

This research is supported by the Canadian Institutes of Health Research (FDN-148415, CHRPJ 523597-18) and the Natural Sciences and Engineering Research Council of Canada (2016-06090, CHRPJ 523597-18).

### Authors contributions

J.D., S.G., E.H.S. and S.O.K. conceived the experiments. J.D. and S.G. designed and performed the experiments. S.G. conceived the theoretical model. J.C. fabricated the chips. J. D. and A. M. fabricated electrodes. H. Y. and W. Z. performed conjugation of ferrocence to oligo and antibody to oligo. S. A., J.D., and S.G. performed experiments with animal model and J.D., S.G., E.H.S. and S.O.K. co-wrote the paper.

### Competing interests

We declare no competing interests.

## References

1. Kim, J., Campbell, A. S., de Ávila, B. E. F. & Wang, J. Wearable biosensors for healthcare monitoring. Nat. Biotechnol. 37, 389–406 (2019).

2. Bandodkar, A. J., Jeang, W. J., Ghaffari, R. & Rogers, J. A. wearable sensors for biochemical sweat analysis. Annu. Rev. Anal. Chem. 12, 1–22 (2019).

3. Giljohann, D. A. & Mirkin, C. A. Drivers of biodiagnostic development. Nature 462, 461–464 (2009).

4. Gaster, R. S. et al. Matrix-insensitive protein assays push the limits of biosensors in medicine. Nat. Med. 15, 1327–1332 (2009).

5. Rong, G., Corrie, S. R. & Clark, H. A. In vivo biosensing: progress and perspectives. ACS Sensors 2, 327–338 (2017).

6. Gao, W. et al. Fully integrated wearable sensor arrays for multiplexed in situ perspiration analysis. Nature 529, 509–514 (2016).

7. Wassum, K. M. et al. Transient extracellular glutamate events in the basolateral amygdala track reward-seeking actions. J. Neurosci. 32, 2734–2746 (2012).

8. Sarter, M. & Kim, Y. Interpreting chemical neurotransmission in vivo: techniques, time scales, and theories. ACS Chem. Neurosci. 6, 8–10 (2015).

9. Lipani, L. et al. Non-invasive, transdermal, path-selective and specific glucose monitoring via a graphene-based platform. Nat. Nanotechnol. 13, 504–511 (2018).

10. Lee, H. et al. A graphene-based electrochemical device with thermoresponsive microneedles for diabetes monitoring and therapy. Nat. Nanotechnol. 11, 566–572 (2016).

11. Bindra, D. S. et al. Design and in vitro studies of a needle-type glucose sensor for subcutaneous monitoring. Anal. Chem. 63, 1692–1696 (1991).

12. Koh, A. et al. A soft, wearable microfluidic device for the capture, storage, and colorimetric sensing of sweat. Sci. Transl. Med. 8, 366ra165 (2016).

13. Kang, D. et al. New architecture for reagentless, protein-based electrochemical biosensors. J. Am. Chem. Soc. 139, 12113–12116 (2017).

14. Arroyo-Currás, N. et al. Real-time measurement of small molecules directly in awake, ambulatory animals. Proc. Natl. Acad. Sci. U. S. A. 114, 645–650 (2017).

15. Ferguson, B. S. et al. Real-time, aptamer-based tracking of circulating therapeutic agents in living animals. Sci. Transl. Med. 5, 213ra165 (2013).

16. Mage, P. L. et al. Closed-loop control of circulating drug levels in live animals. Nat. Biomed. Eng. 1, 1–10 (2017).

17. Somasundaram, S. & Easley, C. J. A nucleic acid nanostructure built through on-electrode ligation for electrochemical detection of a broad range of analytes. J. Am. Chem. Soc. 141, 11721–11726 (2019).

18. Nakatsuka, N. et al. Aptamer–field-effect transistors overcome Debye length limitations for small-molecule sensing. Science 362, 319–324 (2018).

19. Huang, K. C. & White, R. J. Random walk on a leash: a simple single-molecule diffusion model for surface-tethered redox molecules with flexible linkers. J. Am. Chem. Soc. 135, 12808–12817 (2013).

20. Kelley, S. O. et al. Orienting DNA helices on gold using applied electric fields. Langmuir 14, 6781–6784 (1998).

21. Erdmann, M., David, R., Fornof, A. & Gaub, H. E. Electrically controlled DNA adhesion. Nat. Nanotechnol. 5, 154–159 (2010).

22. Langer, A., Kaiser, W., Svejda, M., Schwertler, P. & Rant, U. Molecular dynamics of DNA-protein conjugates on electrified surfaces: solutions to the drift-diffusion equation. J. Phys. Chem. B 118, 597–607 (2014).

23. Herman, E. H. & Ferrans, V. J. Preclinical animal models of cardiac protection from anthracycline-induced cardiotoxicity. Semin. Oncol. 25, 15–21 (1998).

24. Yang, Y. et al. A laser-engraved wearable sensor for sensitive detection of uric acid and tyrosine in sweat. Nat. Biotechnol. 38, 217–224 (2020).

25. Xu, S. et al. Soft microfluidic assemblies of sensors, circuits, and radios for the skin. Science 344, 70–74 (2014).

26. Wang, S. et al. Skin electronics from scalable fabrication of an intrinsically stretchable transistor array. Nature 555, 83–88 (2018).

